# Analysis of the influence of gradual changes in matrix sentence similarity on neural envelope tracking

**DOI:** 10.64898/2026.09.21.752993

**Authors:** Till Habersetzer, Bernd T. Meyer, Andreas Radeloff

**Author notes:** Corresponding author, Communication Acoustics, Carl von Ossietzky Universität Oldenburg, Ammerländer Heerstraße 114-118 26129 Oldenburg, Germany.

## Abstract

Neural tracking of speech is a well-established phenomenon in neuroscience. However, for speech signals with a fixed structure, significant correlations between speech envelopes and neurophysiological representations occur even for unheard sentences. We exploit a structured speech-in-noise matrix hearing test (Oldenburger Sentence Test, OLSA) to systematically quantify the relationship between acoustic sentence similarity and neural tracking. Simultaneous magnetoencephalography (MEG) and 76-channel electroencephalography (EEG) data, including 16 channels positioned directly around the ears (ear-EEG), were recorded from 21 young adults with normal hearing during the presentation of clean-speech audiobooks and OLSA sentences at six signal-to-noise ratios. A linear decoder trained on audiobooks reconstructed OLSA sentence envelopes. Reconstruction accuracies were compared using a linear mixed model across heard (matched) and unheard (mismatched) sentences of varying acoustic similarity. Significant reconstruction accuracies were achieved across MEG, EEG, and ear-EEG for both matched and mismatched sentences. For mismatched sentences, these accuracies gradually increased with their acoustic similarity to the heard speech data. The high similarity between sentences, which is especially prominent in matrix tests, can cause significant spurious tracking for mismatched stimuli. This effect can reach levels comparable to those of matched sentences and can be mistaken for true neural tracking. Robust neural tracking across modalities further supported the established viability of ear-EEG compared to whole-head systems.

## Introduction

Neural speech tracking is a well-established phenomenon in neuroscience in which neurophysiological responses time-lock to specific features in a continuous speech signal (Brodbeck & Simon, 2020; Obleser & Kayser, 2019). Investigating the tracking of various acoustic and linguistic features provides an objective and ecologically valid method for studying auditory and language processes in the brain during natural speech perception (Gillis et al., 2022).

Extensive research has been dedicated to evaluating the link between neural speech tracking and speech intelligibility. A particularly prominent feature used to investigate this is the speech envelope, as its low temporal modulation frequencies are crucial for speech comprehension (Elliott & Theunissen, 2009; Peelle & Davis, 2012; Shannon et al., 1995). However, increasing evidence suggests that neural envelope tracking is a necessary prerequisite, rather than a sufficient condition, for intelligibility (Gillis et al., 2022). Because tracking occurs even in response to unintelligible signals, such as foreign languages (Etard & Reichenbach, 2019), it does not guarantee actual comprehension.

A widely adopted framework for computing neural tracking is based on the Temporal Response Function (TRF) (Crosse et al., 2016). This approach establishes a linear mapping between stimulus features (e.g., the speech envelope) and brain signals. It can be implemented either as a forward model (encoder) that predicts brain signals from speech features or as a backward model (decoder) that reconstructs speech features from the neural responses. The accuracy of this mapping, typically measured using a correlation coefficient, quantifies neural tracking as the linear dependency between speech and brain signals, reflecting the extent to which specific features are represented in the neural response (Brodbeck et al., 2018).

For a decoder, the reconstruction accuracy of the presented speech, serving as a measure of neural tracking, is calculated as the correlation between the reconstructed and original envelopes of the speech presented during the neural recording. To assess statistical significance, a null distribution is generated to estimate the correlations that can arise purely by chance in the absence of a true relationship between brain signals and speech features (Crosse et al., 2021). This is typically achieved by repeatedly correlating the reconstructed envelopes with mismatched speech envelopes, thus establishing a chance-level baseline.

Comparably large correlations between the reconstructions and the speech envelopes can be observed even for signals that have not been presented to the listener which can be mistaken for genuine neural tracking. This effect becomes more pronounced when the original and mismatched sentences are highly similar. An example of a speech material with such similarity between sentences is the matrix test, first developed by Hagerman (1982). Matrix tests are widely used in speech audiometry to assess intelligibility and evaluate auditory function in both research and clinical practice. These tests contain target sentences with a fixed structure (name, verb, numeral, adjective and object, e.g., “Peter sees eight wet stones”) taken from a word matrix, where each column represents a word category with 10 alternatives. A random sentence is constructed by choosing a random word from each word category in this order. This design provides a highly reliable and repeatable speech-in-noise measurement that works across languages (Akeroyd et al., 2015; Brand & Wagener, 2017; Kollmeier et al., 2015).

Therefore, these matrix tests have also been explored for objective speech audiometry by directly measuring cortical activity through neural tracking (Lesenfants et al., 2019; Muncke et al., 2022; Vanthornhout et al., 2018).

Exploiting the rigid syntactic structure of matrix tests, which yields natural speech with similar envelopes, this study quantifies the mismatch between sentence envelopes and its impact on estimated reconstruction accuracies. By gradually varying the degree of this mismatch, either by completely exchanging or shifting the compared envelopes, we systematically explore the expected range of reconstruction accuracies. To do so, we analyze a dataset (Habersetzer et al., 2026) that employed a German matrix test (the Oldenburger Satztest, OLSA) (Wagener, Brand, & Kollmeier, 1999a, 1999b; Wagener, Kühnel, & Kollmeier, 1999) for objective speech assessment using magnetoencephalography (MEG), whole-head electroencephalography (EEG) and an around ear-EEG. The dataset contains neural responses to OLSA sentences presented at various signal-to-noise ratios (SNRs), as well as to clean speech from audiobooks. This multimodal dataset enables us to evaluate the consistency of reconstruction accuracies across different sensor configurations. Furthermore, comparing whole-head systems to ear-EEG allows us to assess whether neural tracking characteristics remain consistent when transitioning from controlled laboratory setups to more mobile, potentially clinically applicable frameworks.

Specifically, we aim to explicitly investigate matrix tests to (i) characterize the statistical relationship between completely mismatched or shifted sentences with varying envelope similarity and its impact on estimated reconstruction accuracies, and (ii) highlight the range of chance-level correlations expected for acoustically similar but mismatched (unheard) speech. Finally, we (iii) test the hypothesis that this effect is robustly observed across whole-head MEG, cap-EEG and sparser ear-EEG, with overall correlation magnitudes decreasing in that order.

## Materials and Methods

### Participants

Twenty-one native German speakers (12 female, 9 male; mean age 27 ± 4 years) participated in the study. All participants were recruited from the university environment and reported no history of neurological disorders. Inclusion required normal or corrected-to-normal vision and normal hearing. The latter was confirmed through a pure-tone audiogram, with an inclusion threshold of ≤ 20 dB HL for the four-frequency pure-tone average (4fPTA: 0.5, 1, 2, 4 kHz) in the better ear (Humes, 2019). Handedness was not used as an exclusion criterion. All participants provided their informed written consent, including permission for the open sharing of anonymized data and were financially compensated. The protocol was approved by the University Ethics Committee (Ref: EK/2021/181) and adhered to the Declaration of Helsinki.

### Experimental Paradigm

The study comprised three sessions per participant: a screening session (ses-00) followed by two recording sessions (ses-01 and ses-02). Data collection for the cohort spanned four months. While the protocol included structural MRI scans and MEG/EEG recordings of transient stimuli such as clicks, these data fall outside the scope of the current research questions and are excluded from the present analysis.

The first session (ses-00) served to screen participants and familiarize them with the MEG environment. Following a standard pure-tone audiogram in a sound-attenuated booth, participants were positioned in the MEG to assess comfort, head-size compatibility and potential artifacts. In this session, as well as at the beginning of each subsequent recording session (ses-01, ses-02), participants completed two runs of speech audiometry inside the MEG to determine their individual behavioral speech reception threshold (SRT), defined as the signal-to-noise ratio yielding a 50 % word recognition rate. The two main recording sessions focused on simultaneous MEG and EEG acquisition and were structurally identical. Each session began with participant preparation (≈ 1 h), followed by data collection inside the magnetically shielded room (≈ 2 h).

### Tasks and Stimuli

#### Behavioral Assessment

##### Audiogram

Pure-tone air conduction thresholds were determined for each ear independently. Measurements were obtained using an Oscilla A50 audiometer (Oscilla, Aarhus, Denmark) in a sound-attenuated booth, covering a frequency range of 0.125, 0.25, 0.5, 0.75, 1, 1.5, 2, 3, 4, 6 and 8 kHz.

##### OLSA

Speech intelligibility was assessed using the German OLSA matrix test. We used open-access synthesized female speech, which has been shown to yield SRTs comparable to those of natural speech (Nuesse et al., 2019). Sentences are constructed from a 50-word matrix (10 alternatives across five syntactic positions: name, verb, numeral, adjective and object) and presented in phonetically balanced lists of 20 sentences to ensure comparable intelligibility. The stimuli were embedded in continuous, stationary speech-shaped noise (fixed at 65 dB SPL) sharing the same long-term spectrum as the speech material. The SRT was determined using an adaptive procedure in which participants repeated recognized words via the MEG intercom to the test supervisor. Measurements were conducted inside the magnetically shielded room using insert earphones. Six measurements (two per session) were obtained for each subject. The task was controlled via a custom Python implementation, which will be made openly available upon publication.

#### Experimental Tasks

Simultaneous MEG and EEG recordings were acquired during two tasks: Listening to audiobooks and to OLSA sentences. The block sequence was fixed, comprising two OLSA blocks, followed by two audiobook blocks and a final OLSA block.

##### Audiobook

Stimuli comprised two German public-domain short stories: “Das schwatzende Herz” (“The Tell-Tale Heart”) by E.A. Poe and “Der gestohlene Bazillus” (“The Stolen Bacillus”) by H.G. Wells. Clean audio was synthesized via a text-to-speech engine (ElevenLabs) using a custom female voice model based on a female speaker from the research group and presented at a long-term average RMS level of 65 dB SPL. Each story lasted approximately 16 min, divided into two runs of equal length. To ensure vigilance, participants answered three comprehension questions following each run. The presentation order was fixed (ses-01: Poe; ses-02: Wells) and participants were instructed to maintain fixation on a central cross throughout the task to minimize eye movements.

##### OLSA

Each recording session (ses-01, ses-02) comprised 540 OLSA sentences, distributed across three runs (200, 200 and 140 sentences). The OLSA speech material was restricted to 100 unique sentences derived from five fixed, 20-sentence lists. From this pool, 27 list presentations were drawn and mapped to six target intelligibility levels (0 %, 20 %, 40 %, 50 %, 80 % and 100 %), which were held constant across all participants. To ensure balanced exposure, each of the five base lists was paired exactly once with each of the five highest intelligibility levels (20 % to 100 %). This arrangement yielded exactly 100 sentences per SNR condition for these upper levels (25 lists total). The 0 % condition was added after piloting to incorporate a performance floor, but it was restricted to only 40 sentences to avoid excessively prolonging the experiment. The presentation order of all list-SNR combinations was randomized across the session, with the constraint that the first list of every run had an intelligibility ≥ 50 % to ensure engagement and signal onset awareness. Throughout the task, background noise was kept constant at 65 dB SPL, while the speech level was varied. Task engagement was monitored via a 2-alternative forced-choice paradigm using a button box. A response was mandatory after every list, with an additional 60 % probability of an intermediate response occurring between sentences 6 and 14 (position sampled from N (10, 4)) to maintain participant engagement. Participants selected the correct target word from two options within 4 s, followed by immediate visual feedback. Sentences were separated by a jittered inter-stimulus interval (0.8 s to 1.2 s) and initial trial onset was jittered between 1 s and 1.5 s. Target intelligibilities (*p*) were mapped to individual SNRs (*x*) using the individually fitted psychometric function (Brand & Kollmeier, 2002):

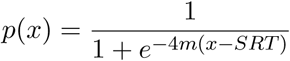

where the slope *m* was fixed at 0.13 (Nuesse et al., 2019) and the *SRT* was determined individually from the fourth behavioral OLSA measurement (ses-01, run-02) to ensure performance stability (Nuesse et al., 2019; Wagener, Brand, & Kollmeier, 1999b). Participants therefore completed three training lists over two days prior to MEG/EEG data collection. Mapping fixed intelligibility levels to individualized SNRs, rather than utilizing uniform SNRs across all subjects, was chosen to optimize sampling along the steep incline of individual psychometric curves. Individualized SNRs were derived as follows: fixed anchor points were used for the floor (≈0 % at −40 dB) and ceiling (≈100 % at 0 dB) conditions, while intermediate levels were anchored to individual SRTs. This mapping yielded the following target SNRs: 20 % (SRT − 2.7 dB), 40 % (SRT − 0.8 dB), 50 % (SRT) and 80 % (SRT + 2.7 dB). Prior to the task, participants completed a two-list training run (40 sentences) and were instructed to maintain fixation on a central cross on the screen.

### Data Acquisition

#### MEG and EEG

Neurophysiological data were recorded in a magnetically shielded room (Vacuumschmelze, Hanau, Germany) using a 306-channel Elekta Neuromag Triux system (Elekta Oy, Helsinki, Finland), comprising 102 magnetometers and 204 planar gradiometers. Subjects were seated in an upright position (68*^◦^*). Simultaneous EEG was acquired via a custom 76-channel cap (BC-MEG-76-X2; Easycap, Wörthsee, Germany) that augmented the standard Elekta 64-channel layout with 12 additional around-ear electrodes (6 per side) to enhance coverage of temporal regions (see Figure 1). The EEG setup utilized a right-nasal reference electrode, a forehead ground electrode and maintained impedances below 5 kΩ whenever possible. Continuous head position tracking was performed using five HPI coils. During preparation, the positions of the three anatomical landmarks, HPI coils, EEG electrodes and *>* 200 head-shape points were digitized using a Polhemus Fastrak device (Polhemus, Colchester, VT, USA). Physiological artifacts were monitored via one electrocardiogram (ECG) and two bipolar electrooculography (EOG) channels (vertical: VEOG, horizontal: HEOG). All signals were sampled at 1000 Hz with an online band-pass filter of 0.1 Hz to 330 Hz. To ensure consistent head position and sensor proximity, participants repositioned their heads against the top of the helmet prior to each recording run.

**Figure 1:**
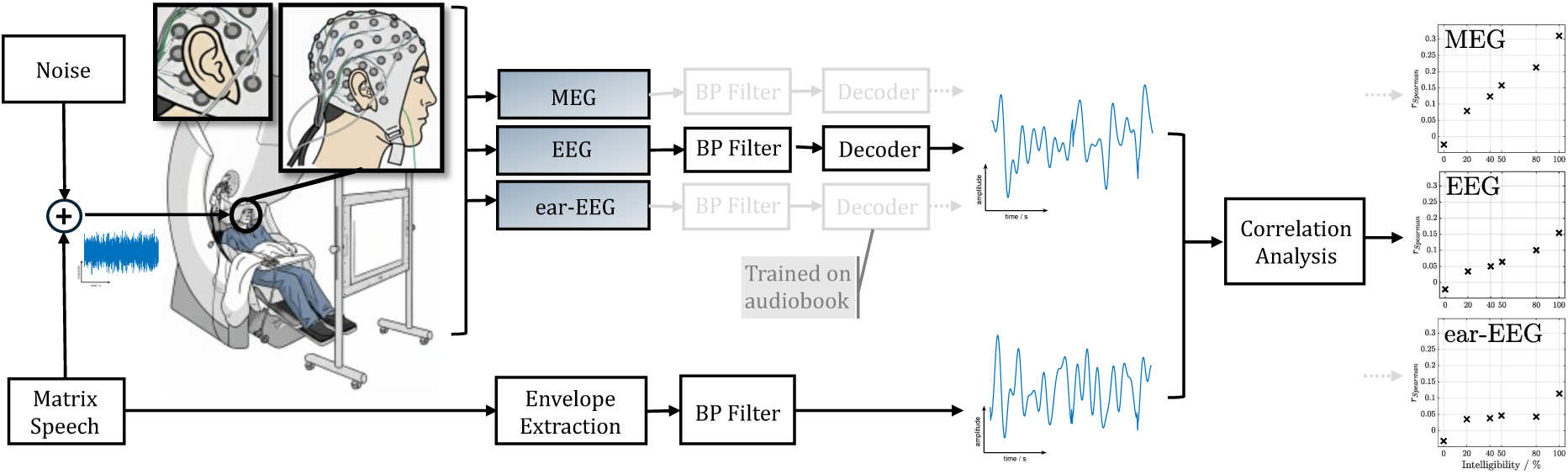
Overview of the experimental setup and analysis pipeline. Participants were seated in the MEG scanner while wearing a high-density EEG-cap featuring a subset of electrodes positioned around the ears (termed ear-EEG). MEG and EEG were acquired simultaneously. For the analysis, we distinguish between three sensor modalities: MEG, EEG (full EEG-cap including all ear channels) and ear-EEG. Acoustic stimulation was delivered via insert earphones and consisted of OLSA sentences presented at varying SNRs corresponding to fixed intelligibility levels. For the data analysis, recordings from each modality were bandpass filtered and a linear decoder trained on clean-speech audiobooks was used to reconstruct the OLSA speech envelopes. For evaluation, the clean-speech OLSA envelopes were extracted from the audio files and filtered identically. Neural envelope tracking was quantified by calculating the correlation between the reconstructed and true envelopes. The figure concept is adapted from Vanthornhout et al. (2018).

#### Hardware, Software and Synchronization

Diotic audio stimuli were presented at 44.1 kHz using insert earphones (CareFusion Model TIP 300, 300 Ω) driven by a TDT HB7 headphone driver (Tucker-Davis Technologies, Alachua, FL, USA) and an RME Fireface UCX soundcard (RME, Haimhausen, Germany). Acoustic calibration was conducted via a Brüel & Kjær (B&K) Type 4157 Ear Simulator with a DB-2012 Ear Mould Adaptor, a Type 2669 microphone preamplifier and a Type 2610 measuring amplifier calibrated by a B&K Type 4231 Sound Calibrator. Speech-shaped OLSA noise served as the calibration reference for behavioral audiometry, the MEG OLSA and audiobook sound levels. Visual stimuli were projected via a PROPixx MRI/MEG projector system (VPixx Technologies, Saint-Bruno-de-Montarville, QC, Canada), with responses collected using a fiber-optic button box (RESPONSEPixx/MRI, VPixx Technologies) and a DATAPixx3 hub (VPixx Technologies). The experimental framework was implemented in Python 3.8 and PsychoPy (v2024.1.4) (Peirce et al., 2019), while audio delivery was handled via the SoundMexPro engine (Berg, 2024) to ensure low-latency playback. Temporal synchronization was achieved via a custom FPGA-based trigger box that converted the soundcard’s SPDIF output into 5 V TTL triggers for the MEG. These triggers indicated precise sentence onsets for the OLSA task and 1 s intervals for the audiobook task.

### Data Analysis

Data analysis was performed for all 21 participants and is illustrated in Figure 1. Any subject-specific exclusions and protocol deviations are reported in Table S1 in the supplementary material. Because the same measurement setup was used across sessions, data from both recording sessions (ses-01 and ses-02) were pooled to maximize statistical power and streamline the analysis. Consequently, the data for each participant included four audiobook runs, comprising two audiobooks each split into two runs, providing approximately 32 minutes of continuous speech. Additionally, the data included six OLSA runs totaling 1080 sentences. This consisted of 200 sentences per SNR (intelligibility), with the exception of the −40 dB condition, which contained only 80 sentences.

#### Signal Processing and Preprocessing

Neural MEG recordings were preprocessed in MNE-Python (Gramfort et al., 2013) using Maxwell filtering (Taulu & Kajola, 2005; Taulu & Simola, 2006), which incorporated spatiotemporal signal space separation (tSSS), head movement correction and transformation to a common head position (referenced to the *ses-01_task-olsa_run-02* recording) to ensure consistent sensor-to-head alignment across recordings. The PyPREP Python library (Appelhoff et al., 2025; Bigdely-Shamlo et al., 2015) was used to automatically detect bad EEG channels. Subsequent analysis was conducted in MATLAB using the FieldTrip toolbox (Oostenveld et al., 2011). Both MEG and EEG signals were filtered between 0.5 Hz to 8 Hz (delta and theta bands) using a windowed-sinc finite impulse response filter (firws) (Widmann et al., 2015). Three sensor modalities were defined for the analysis: (1) **MEG**, including all 306 channels (magnetometers and gradiometers), (2) **EEG**, comprising all 76 channels (standard 64-channel layout plus 12 around-ear electrodes) re-referenced to a common average and (3) **ear-EEG**, a 16-channel subset utilizing the 12 around-ear electrodes and channels T7, TP7, T8 and TP8 from the 64-channel layout (8 channels per ear). See Figure S1 in the supplemental material for details on the EEG and ear-EEG layouts. To ensure robustness and model galvanic separation, the ear-EEG data were re-referenced separately for each ear using the local average potential of that ear’s respective channels. Prior to referencing, the detected bad EEG channels were interpolated using spline interpolation, performed separately for the full EEG and ear-EEG layout. Audiobook data were segmented into continuous 60 s epochs, whereas OLSA recordings were epoched per sentence from stimulus onset to 0.5 s post-stimulus. Finally, signals for both tasks were downsampled to 64 Hz. The recorded EOG and ECG data were not utilized for artifact removal, as the linear decoding approach appeared to be sufficiently robust without additional corrections. Speech envelopes were extracted following Biesmans et al. (2016) using a 28-channel Gammatone filterbank, with channels spaced by one equivalent rectangular bandwidth between 5 Hz and 5000 Hz. The absolute output of each channel was raised to the power of 0.6 and averaged across channels to derive a broadband envelope. Subsequently, this envelope was filtered between 0.5 Hz to 8 Hz and downsampled to 64 Hz to match the neural data.

#### Decoder Training

Speech envelopes were reconstructed from multivariate neural recordings using a backward linear model (decoder) with integration latencies of 0 ms to 400 ms, implemented via the mTRF-Toolbox (Crosse et al., 2016, 2021) in MATLAB. Before training, the audio envelopes were scaled in a range of [−1, 1] via peak amplitude scaling. Multichannel MEG and EEG data were *z*-scored globally across all trials and sensors, rather than channel-by-channel, to preserve relative power across channels. Therefore, a global mean and standard deviation were calculated independently for the EEG, magnetometer and gradiometer sensor groups to account for their different physical units and magnitudes. For each subject, the decoder was trained using pooled audiobook data from both sessions (approximately 32 epochs, each 60 s). First, the regularization parameter (*λ*) was optimized across all audiobook trials using leave-one-out cross-validation. Following this optimization, the final model was trained on this complete set of trials using the selected *λ*.

#### Decoder Application and Evaluation

Neural envelope tracking was quantified by applying the subject-specific decoders to the strictly held-out OLSA recordings for each sensor modality and intelligibility level. For each sentence, an envelope was reconstructed (i.e., predicted) and paired with its corresponding acoustic envelope. To ensure temporal alignment, valid speech segments were cropped to a shared minimum duration prior to concatenation into continuous vectors (comprising 200 sentences or 80 for the −40 dB condition). Reconstruction accuracy was determined using Spearman correlations between predicted and acoustic envelopes for four control conditions:

- **Matched Sentences:** The reconstructed envelopes were correlated with the vector of matched acoustic envelopes, representing the neural tracking measure.
- **Null Distribution:** To eliminate sentence matching and temporal alignment, a distribution was generated via 1000 iterations. In each iteration, sentence pairings were randomized as a derangement (ensuring no true pairs) and the acoustic envelope for each pair was randomly circularly shifted by at least 1 s. The reconstructed and acoustic envelopes were first separately concatenated and then correlated with each other. The final reconstruction accuracy is defined as the mean correlation across all iterations.
- **Shuffled Sentences:** Generated identically to the Null Distribution (using 1000 deranged pairings), but without the circular shift, isolating the contribution of sentence-level acoustic structure shared across the corpus.
- **Mean-Sentence:** Reconstructed envelopes were correlated with a concatenated vector of the mean envelope (the average temporal envelope across all sentence envelopes in the corpus) to isolate the general rhythmic structure shared across the corpus.

Because this study predominantly evaluates mismatched control audio, we use *reconstruction accuracy* as our overarching term, even though it is slightly misleading to speak of reconstruction when comparing against a mismatched envelope. We reserve *neural tracking* for instances where the reconstructed envelope is compared against the actually heard audio, as this specifically measures stimulus tracking. For all mismatched conditions, we refer to *reconstruction accuracies*, as these evaluate audio that was not actively tracked.

All correlation coefficients were Fisher *z*-transformed prior to averaging and subsequently back-transformed. To quantify the increase in reconstruction accuracy with speech intelligibility, we employed linear mixed-effects models (LMMs). The model was defined in matrix form as:

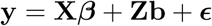

In this formulation, **y** is the vector of reconstruction accuracies, while **X** and **Z** are the design matrices for fixed and random effects, respectively. The vector ***β*** contains the global fixed-effect coefficients for the intercept (*β*_0_) and the slope (*β*_1_), while **b** represents the subject-specific random effects for the intercept (*b*_0_*_,s_*) and the slope (*b*_1_*_,s_*). Finally, ***ϵ*** denotes the residual error. By considering the observation for a single subject *s* at a given intelligibility level *i* as a single row within this matrix equation, the model is expressed as:

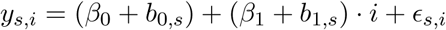

The fixed-effect slope (*β*_1_), hereafter denoted as *β_I_*, captures the group-level rate at which reconstruction accuracy improves with higher intelligibility. For numerical stability, intelligibility levels were expressed as proportions (0 to 1) rather than percentages. Consequently, *β_I_* reflects a dimensionless rate of change in reconstruction accuracy per unit of intelligibility. This normalization allows a positive slope to be intuitively interpreted as the group’s maximum reconstruction accuracy. Given that intelligibility scales from zero to one, assuming a reconstruction accuracy near zero at 0 % intelligibility means that *β_I_* directly approximates the expected accuracy at 100 % intelligibility.

We further investigated the relationship between reconstruction accuracies and the acoustic similarity among the OLSA sentences. To assess acoustic similarity in the Shuffled Sentences condition, we calculated the Spearman correlation across all unique envelope pairs (excluding identical pairs). For the Mean-Sentence condition, we correlated the global mean envelope with each individual OLSA envelope. For both conditions, the final acoustic similarity metric was defined as the mean of these correlation values. Beyond these controls, we performed a more granular analysis to evaluate how specific acoustic properties and temporal alignments influence reconstruction accuracy. Specifically, we systematically assessed the similarity between individual OLSA envelopes and their time-shifted versions to determine how this similarity impacts reconstruction performance.

##### Inter-Sentence Similarity

To quantify how closely each OLSA sentence envelope resembles the rest of the speech material (i.e., the OLSA corpus), we calculated a mean similarity score (*ρ_mean_*) for each sentence. This metric represents the average Spearman correlation between a given sentence envelope and all other envelopes in the corpus (excluding itself), with each pair cropped to a common minimum length (see Figure 4(a) and (b)). Sentence-specific reconstruction accuracies were computed by correlating reconstructed envelopes with a concatenated vector of that specific sentence’s envelope for each intelligibility level, analogous to the Mean-Sentence condition. An LMM was then fitted to these values across all subjects to estimate a sentence-specific fixed-effect slope (*β_I_*). This procedure is illustrated in Figure 2(a) and (b). Finally, the relationship between acoustic similarity and reconstruction accuracy was evaluated by correlating the *ρ_mean_* scores with their respective neural slopes (*β_I_*). Additionally, we assessed whether sentence-specific reconstruction accuracies differed significantly from the null distribution. For each sentence, a group-level comparison was performed across all subjects using paired t-tests, comparing reconstruction accuracies (*ρ_Spearman_*) against the mean values of the null distribution (Fisher *z*-transformed) for each of the six intelligibility levels. The resulting *p*-values were corrected using the False Discovery Rate (FDR). A sentence was categorized as significantly different if the FDR-corrected *p*-values were below the threshold (*p <* 0.05) for at least three of the five higher intelligibility levels (≥ 20%). See Table S2 in the supplementary material for further details regarding the applied FDR corrections.

**Figure 2:**
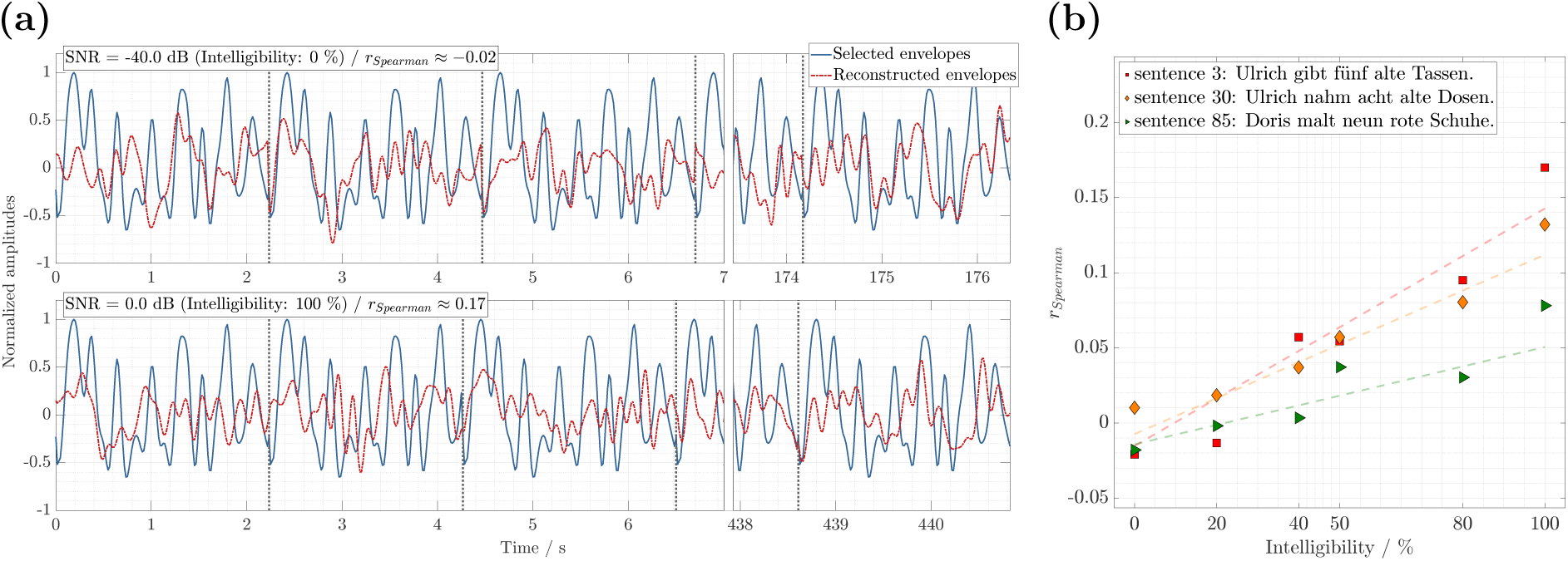
Representative envelope reconstructions and data analysis for subject 10 (MEG, pooled sessions): **(a)** The reconstructed envelope from the decoder (red dashed) is compared to concatenated envelopes of a single sentence (blue solid). As an example, the sentence “Ulrich gibt fünf alte Tassen” is shown for 0 % intelligibility (upper panel) and 100 % intelligibility (lower panel). Correlations were computed by concatenating speech-only segments, comprising 80 sentences for the 0 % condition and 200 sentences for all other levels. This results in the sharp transitions visible between segments. Sentence boundaries are indicated by vertical dashed gray lines. Due to the total signal length, only the initial 7 s and final 3 s are displayed. **(b)** Spearman correlation coefficients as a function of intelligibility. The values derived for the sentence in panel (a) (red squares) are shown alongside those of two additional exemplary sentences (orange diamonds and green triangles), illustrating similar trends. Sentences are ordered according to the fixed-effect slope derived from the LMM. The numbers in the legend indicate the slope rank in descending order (i.e., lower numbers indicate steeper slopes). While this analysis was performed for all OLSA sentences, the dashed lines represent the specific LMM fits for these three examples for a single subject (incorporating the subject-specific random effect intercept and slope). Nonetheless, these curves were derived from a model estimated using data aggregated across all participants.

##### Temporal Alignment and Cross-Correlation

To assess how temporal alignment impacts reconstruction accuracy across various time lags (*τ*) between reconstructed and matched envelopes, a modified cross-correlation function was computed to evaluate shift-specific performance, mirroring the inter-sentence similarity analyses. Utilizing the matched-sentence condition, we also conducted this analysis on concatenated sentence vectors. Within each individual sentence segment, the matched envelope was circularly shifted sample-wise (1 sample ≈ 16 ms) relative to the reconstructed envelope. Following this shift, all segments were concatenated to compute a single overall Spearman correlation coefficient for each intelligibility level. To construct a cross-correlation function centered at zero, the matched envelope was shifted in both positive and negative directions. The maximum allowed lag was restricted to half the duration of the shortest sentence in the corpus (≈ 2 s), constraining the maximum shift to ±1 s. Subsequently, an LMM was fitted to the data for each shift to estimate a fixed-effect slope (*β_I_*(*τ*)). Significant difference against a null distribution at each time-shift was assessed using the same procedure applied for inter-sentence similarity.

To relate these lag-specific reconstruction accuracies to the linguistic content of the OLSA envelopes (such as phoneme or word rates), a comparable cross-correlation function and the temporal modulation spectrum were computed directly from the speech material. This cross-correlation (*ρ_shift_*(*τ*)) addresses the acoustic similarities between OLSA envelopes at corresponding time lags and was calculated across all possible envelope pairs. Two comprehensive vectors were constructed by concatenating all 9900 unique, non-identical sentence envelope pairs (100 × 99, accounting for pairing order and excluding identical pairs). Mirroring the former analysis, the paired envelopes were shifted sample-wise within their respective sentence segments, concatenated across all pairs and correlated at each lag. Because the full permutation matrix of sentence pairings was utilized, this cross-correlation function is symmetric around zero.

Finally, the temporal modulation spectrum of the OLSA envelopes was derived by averaging the individual sentence modulation spectra. Prior to computing the Fast Fourier Transform, all envelopes were tapered with a Tukey window (10 % taper ratio) to minimize edge artifacts while preserving the core time-domain signal. Additionally, word and phoneme rates were calculated for the OLSA sentences by dividing the word or phoneme count of each sentence by its respective duration.

## Results

The following results demonstrate how reconstruction accuracy changes gradually when systematically varying the mismatched or time-shifted matched envelopes. We quantified the increase in accuracy across intelligibilities using fixed-effect slopes and related these measures to the acoustic similarity scores of the underlying envelopes. The analysis was conducted in two stages. First, we assessed reconstruction accuracies across four control conditions: the matched-audio envelope representing the true neural tracking, the mean-sentence envelope, a distribution of shuffled sentences and a null distribution incorporating an additional circular shift. Second, we quantified these changes in more detail by analyzing reconstruction accuracies for individual OLSA sentences and their time-shifted versions.

### Reconstruction Accuracies Across Control Conditions

We evaluated reconstruction accuracies across four control conditions: the Null Distribution (shuffled and circularly shifted), Shuffled Sentences (same but no shift), Mean-Sentence and Matched Sentences. Results for MEG, EEG and ear-EEG are shown in Figure 3, in which reconstruction accuracies (*r_Spearman_*) of all participants are plotted as a function of intelligibility. The performance patterns were consistent across all three sensor modalities. The Null Distribution yielded reconstruction accuracies near zero, confirming the absence of a systematic relationship between the neural signals and the randomly time-shifted mismatched envelopes. In contrast, the remaining three conditions exhibited significant Pearson correlations (*ρ*) between reconstruction accuracy and intelligibility, demonstrating an increase in reconstruction accuracy with speech intelligibility, as also indicated by the significant fixed-effect slopes (*β_I_*) of the fitted LMMs. While the overall trends were similar across sensors, MEG provided the highest reconstruction accuracies and the steepest slopes, followed by EEG and ear-EEG. Across all sensor modalities, the Matched Sentences and Mean-Sentence conditions yielded the highest and most comparable slopes. The Shuffled Sentences condition showed reduced slopes, while the Null Distribution remained flat. Despite these robust group-level trends, substantial inter-subject variability was observed in the individual reconstruction accuracies.

**Figure 3:**
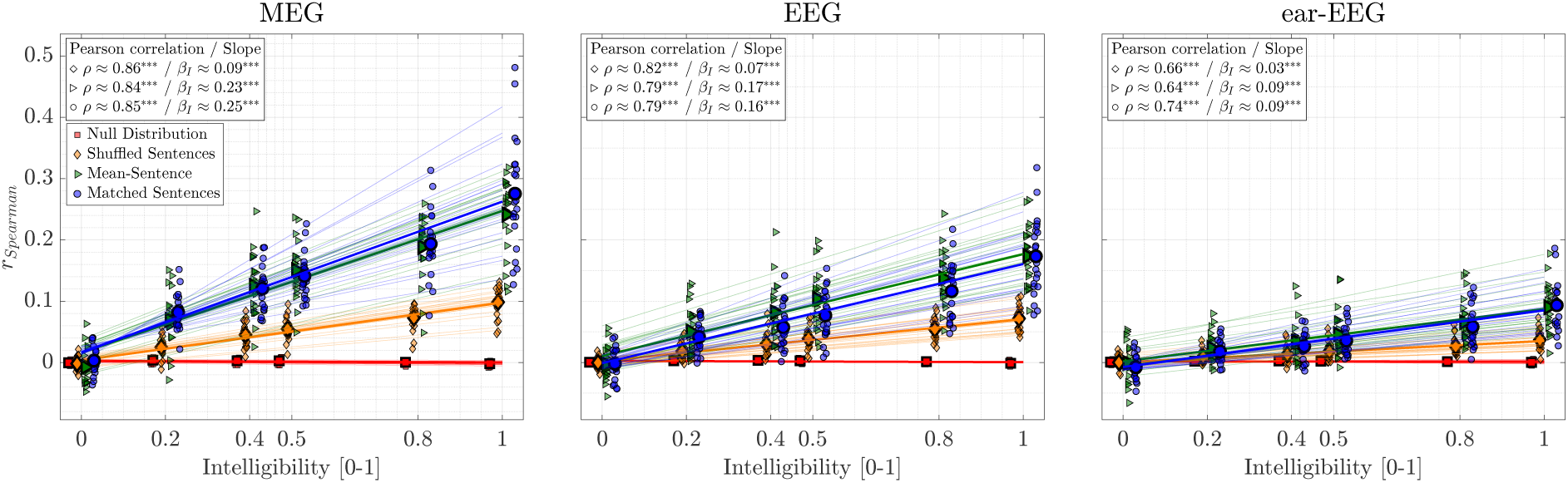
Reconstruction accuracy (*r_Spearman_*) as a function of speech intelligibility for MEG, EEG and ear-EEG. Individual data points are shown alongside grand average means (large symbols) for four conditions: Matched Sentences (blue circles), Mean-Sentence (green triangles), Shuffled Sentences (orange diamonds) and Null Distribution (red squares). To improve visibility, x-axis positions included a horizontal offset between conditions and a slight jitter for individual data points. Thick lines represent LMM fixed-effects, while thin lines represent individual fits. Inset text displays Pearson correlation coefficients (*ρ*) between intelligibility and *r_Spearman_*, alongside fixed-effect slope estimates (*β_I_*). Intelligibility is scaled as a proportion (0–1), yielding a dimensionless slope. All *p*-values were adjusted using FDR. Asterisks indicate significance: *** *p <* 0.001, ** *p <* 0.01, * *p <* 0.05.

### Reconstruction Accuracies and Sentence Similarity

Figure 4(c) depicts the ranked mean similarity scores (*ρ_mean_*) characterizing the inter-sentence similarity for all sentences in the used OLSA corpus. The 100 unique OLSA sentence envelopes exhibit a distinct similarity gradient relative to the rest of the corpus, with Spearman correlation values spanning a range from approximately 0 to 0.25.

**Figure 4:**
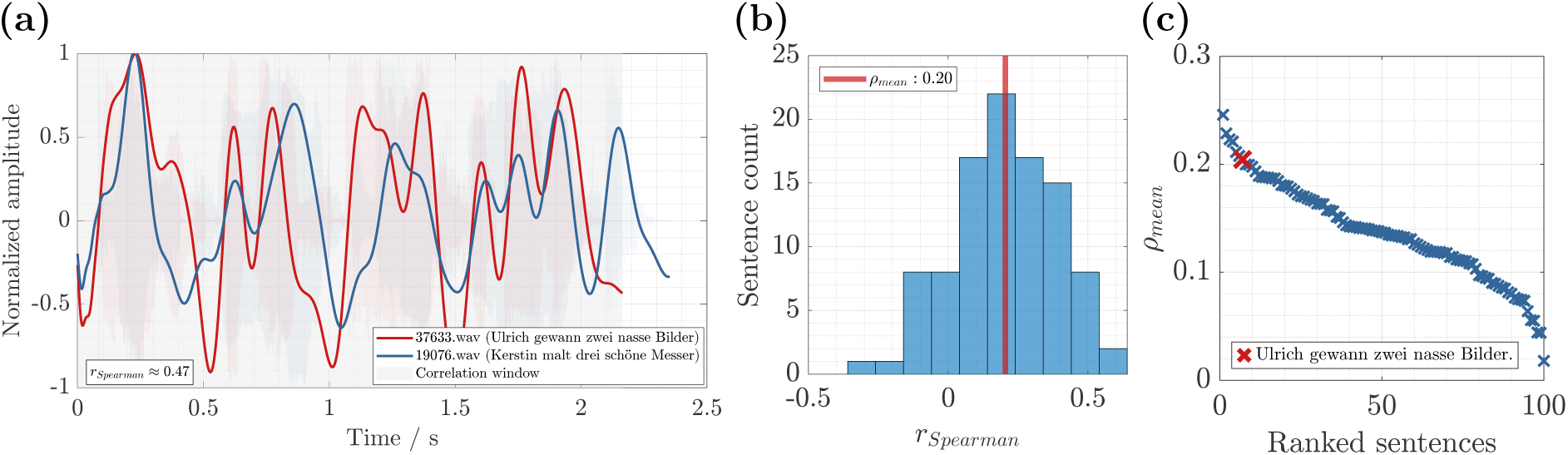
Example of similarity computations between OLSA sentence envelopes. **(a)** Temporal envelopes of two OLSA sentences are shown in blue and red, overlaid on their respective waveforms (background, faded colors). A grey shaded area indicates the time window utilized for calculating the Spearman correlation between the envelopes, with the resulting value displayed in the lower-left inset. **(b)** Distribution of Spearman correlation values (*r_Spearman_*) obtained by correlating the reference sentence (“Ulrich gewann zwei nasse Bilder”) from panel (a) with all other sentences in the OLSA corpus, excluding itself. The distribution mean (*ρ_mean_*) is indicated by a red vertical line and the specific correlation value calculated in panel (a) represents a single data point within this histogram. The number of bins was determined via the square-root choice, 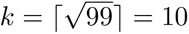. **(c)** Mean correlation values (*ρ_mean_*) for all sentences, calculated according to the procedure in panel (b), are sorted in descending order. The mean value for the sentence in panel (b) is highlighted with a red cross, while the means for all other sentences are indicated by blue crosses.

Figure 5 plots the fixed-effect slopes (*β_I_*) computed for concatenated identical OLSA sentence envelopes against their corresponding mean acoustic similarity scores (*ρ_mean_*). This analysis was conducted across all three modalities (MEG, EEG and ear-EEG) using modality-specific decoders and reconstructions. All three modalities exhibited a significant positive correlation between the mean envelope similarity of the sentences and their respective LMM slopes. MEG yielded the highest slopes, driven by higher reconstruction accuracies across the different intelligibility levels, followed by EEG and ear-EEG, respectively. Furthermore, across all three modalities, most sentences showed reconstruction accuracies that differed significantly from their null distributions and a significant fixed-effect slope. Specifically, these sentence-specific slopes ranged from near zero up to 70 % for MEG and ear-EEG and 80 % for EEG, compared to the maximum slopes achieved with correctly matched sentences. See Figure S3 in the supplementary material for example data and corresponding fits for specific sentences. For a comprehensive comparison, the acoustic similarity and LMM slopes for the Shuffled Sentences and Mean-Sentence conditions are also included. While the Shuffled Sentences condition aligns with the center of the sentence distribution, the Mean-Sentence condition shows markedly higher mean similarity and steeper slopes. Notably, this condition reached approximately 95 %, 105 % and 90 % of the maximum slopes achieved with matched envelopes in MEG, EEG and ear-EEG, respectively (see Figure 3 for comparison).

**Figure 5:**
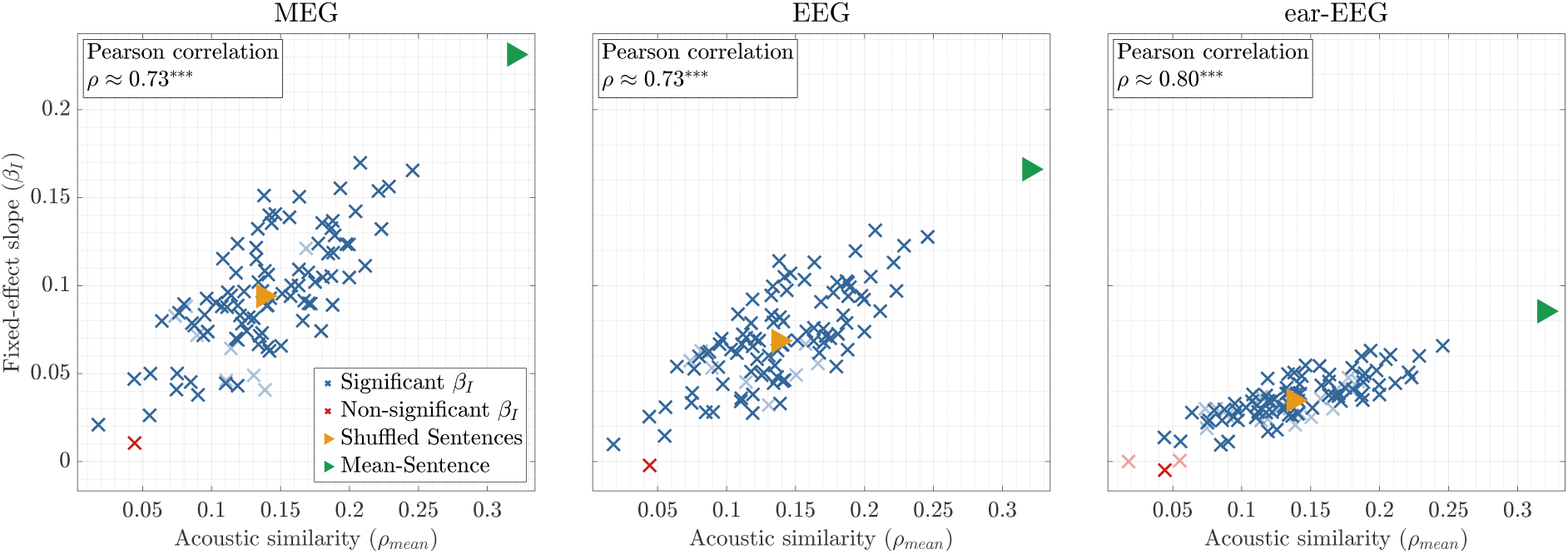
Relationship between sentence-specific mean acoustic similarity and fitted slope. Fixed-effect slopes (*β_I_*) are plotted against mean acoustic envelope similarity (*ρ_mean_*) for individual OLSA sentences (represented by crosses). Blue crosses indicate sentences with a significant fixed-effect slope (different from zero), while red crosses indicate non-significant slopes. Sentences whose reconstruction accuracies did not show a significant difference from their null distributions are marked with faded crosses. Pearson correlation coefficients (*ρ*) between *β_I_* and *ρ_mean_* are provided for MEG, EEG and ear-EEG. Correlations were calculated across all sentences, excluding the Mean-Sentence and Shuffled Sentences conditions. For reference, Mean-Sentence (green triangle) and Shuffled Sentences (orange triangle) conditions are indicated. Asterisks denote significant Pearson correlations (*** *p <* 0.001, ** *p <* 0.01, * *p <* 0.05). All tests were FDR corrected.

### Reconstruction Accuracies and Time-Shift

Figure 6(a) illustrates the fixed-effect slopes (*β_I_*) across circular shifts (*τ*) between the reconstructed and matched envelopes for all three modalities, as well as the grand average curve. While the overall shape of the cross-correlation function is highly consistent across modalities, peak amplitudes decrease systematically from MEG to EEG to ear-EEG. Unlike the symmetric acoustic cross-correlation (Figure 6(b)), which averages bidirectional sentence pairs ((*a, b*) and (*b, a*)), the curve in panel (a) is inherently asymmetric. This asymmetry arises because the computations evaluate a strictly unidirectional relationship (shifting only the matched envelopes) across the concatenated envelope sequences, such that forward and backward circular shifts produce distinct segment overlaps. The primary tracking peak at 0 ms is flanked by alternating positive and negative extrema, reflecting underlying rhythmic periodicities that span a frequency range from 1 Hz to 6.4 Hz. Across all three modalities, reconstruction accuracies differed significantly from their null distributions, and their fixed-effect slopes were significantly different from zero at most local extrema. MEG and EEG demonstrate the broadest temporal clusters of significance, followed by ear-EEG. See Figure S4 in the supplementary material for example data and corresponding fits for specific circular shifts.

**Figure 6:**
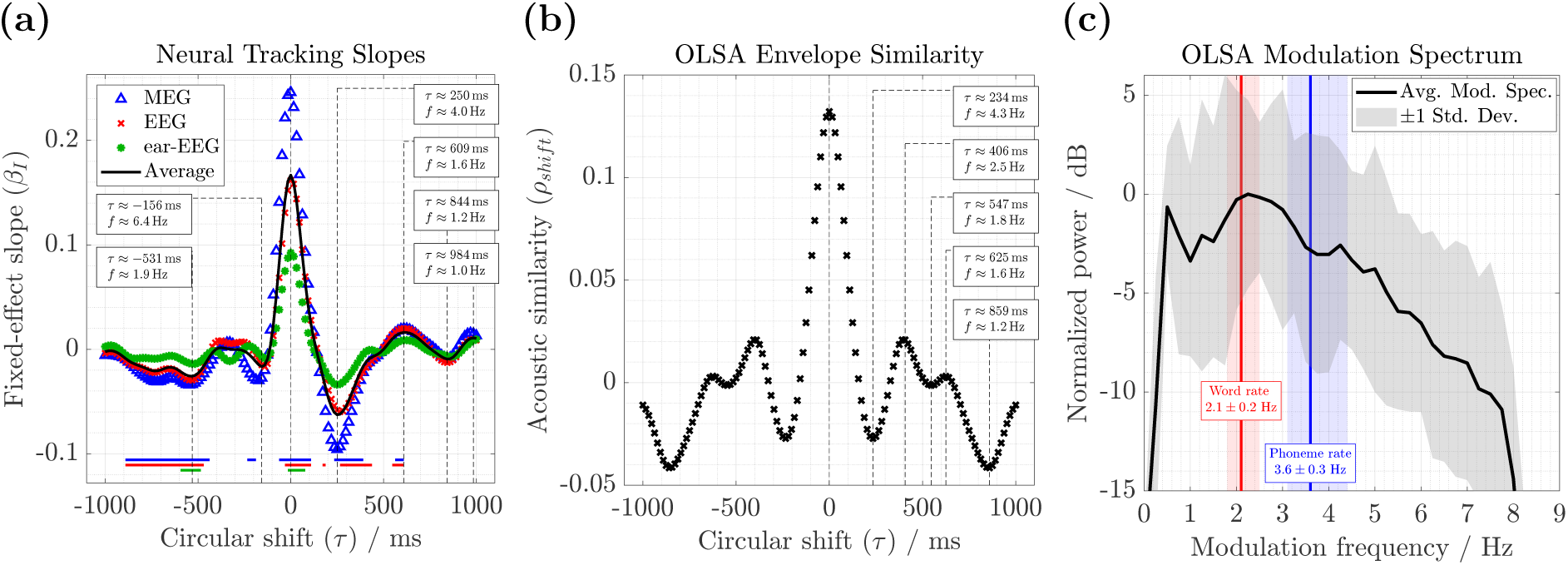
Impact of temporal alignment on reconstruction accuracies and their associated LMM slopes and acoustic envelope similarity. **(a)** Fixed-effect LMM slopes (*β_I_*) for MEG (blue triangles), EEG (red crosses) and ear-EEG (green circles), along with the grand average across modalities (black line), plotted as a function of circular time-shift (*τ*). Models were fitted independently at each lag to quantify the relationship between reconstructed and shifted envelopes for matched sentence pairs across all intelligibility levels. Horizontal bars at the bottom of the axis indicate shift intervals where reconstruction accuracies differed significantly from their null distributions and exhibited significant fixed-effect slopes (both FDR-corrected), with colors denoting the modalities. **(b)** Acoustic similarity (*ρ_shift_*) between linearly shifted concatenated envelope pairs from different OLSA sentences, quantified via Spearman correlation. In Panels (a) and (b), prominent local extrema are highlighted with vertical dashed lines, indicating the shift latency (*τ*) and its corresponding frequency (*f* = 1*/τ*). **(c)** Average temporal modulation spectrum for OLSA sentences (black line, with 1 ± standard deviation indicated by the grey shaded area). The ranges for word rate (red) and phoneme rate (blue), derived from the sentence material, are plotted as background shaded areas bounded by their minimum and maximum values. The mean rates are marked by thick vertical lines and accompanied by text boxes displaying the mean ± standard deviation.

Figure 6(b) depicts the acoustic envelope similarity across time lags (*ρ_shift_*). This curve shares a shape comparable to the cross-correlations in panel (a), particularly along the positive shift axis, with local extrema emerging at latencies corresponding to a frequency range spanning 1.2 Hz to 4.3 Hz. This partially resembles the peak timings observed in panel (a), with the notable exception of the peak near 2.5 Hz.

The average modulation spectrum of the OLSA sentences (derived from the bandpass-filtered envelopes between 0.5 Hz and 8.0 Hz) reveals sustained dominant frequencies up to roughly 5.0 Hz, with a broad peak spanning 2 Hz to 3 Hz. The calculated phoneme and word rates yielded means of 3.6 Hz (range: 3.1 Hz to 4.4 Hz) and 2.1 Hz (range: 1.8 Hz to 2.5 Hz), respectively, aligning directly with the prominent plateau and broad peak in the modulation spectrum.

## Discussion

We observed significant reconstruction accuracies and fitted slopes for both matched and mismatched OLSA sentences, both metrics increasing with speech intelligibility. For mismatched envelopes, the reconstruction accuracies approached true neural tracking values, gradually increasing with acoustic similarity to the OLSA corpus. Furthermore, time-shifted envelopes yielded peaks at lags corresponding to word and phoneme rates. Finally, these findings were consistent across MEG, EEG and ear-EEG.

### Neural Tracking for Matched Sentences

As a baseline result, we observed robust neural tracking using matched predictions and envelopes, with reconstruction accuracies consistently increasing with speech intelligibility (which is linked to the underlying SNR) across all three sensor modalities. This tracking enhancement aligns with extensive previous research that has demonstrated significant correlations between tracking measures and behavioral intelligibility using comparable linear models, particularly within the delta and theta frequency bands (see Ratelle and Tremblay (2025) for a systematic review). EEG remains the most widely adopted modality for these linear approaches (Borges, Alickovic, et al., 2025; Borges, Zaar, et al., 2025b; Cooper et al., 2025; Lesenfants et al., 2019; Vanthornhout et al., 2018), followed by MEG (Ding & Simon, 2013; Habersetzer et al., 2025, 2026; Karunathilake et al., 2023) and ear-EEG (Borges, Alickovic, et al., 2025; Borges, Zaar, et al., 2025a).

### Reconstruction Accuracies for Mismatched Sentences

We observed significant reconstruction accuracies and fitted slopes for mismatched envelope pairs. This mismatch occurred when the evaluated envelope belonged to a different sentence than the presented sentence or when the matched envelope was shifted over time.

#### Inter-Sentence Similarity

To explore mismatched envelopes, we systematically varied the evaluation envelopes and computed sentence-specific reconstruction accuracies and LMM slopes, which were then compared with the acoustic similarity of the sentences to the OLSA speech material. We expected a gradual increase in slope and its correlation with acoustic envelope similarity due to two key factors: (a) the underlying TRF model assumptions and (b) the computational structure of the reconstruction accuracies. (a) The TRF framework assumes the sensory system operates as a linear time invariant (LTI) system, which is approximated by an impulse response function (Crosse et al., 2016). In this study, decoder weights were trained independently on audiobooks to capture general speech properties. Speech envelopes were then reconstructed by convolving these estimated decoder weights with the multivariate brain signals recorded during the OLSA task. Through this convolution, neural responses project their temporal rhythms driven by the stimulus into the envelope reconstructions. Consequently, comparable sentence envelope patterns in the neural responses are preserved in the reconstructions. (b) The computations for sentence-specific LMM slopes and acoustic envelope similarity on both axes are structurally related. When computing the reconstruction accuracy at a given intelligibility level, a given sentence envelope is compared against a broader set of reconstructed envelopes from the OLSA corpus. This comparison set comprises 40 sentences at 0 % intelligibility and 100 sentences for all other intelligibility levels, which doubles when both recording sessions are combined, as done in this analysis. Consequently, reconstruction accuracy reflects how similar a sentence envelope is to the entire set of reconstructed OLSA envelopes. This computational structure closely mirrors the acoustic similarity metric, which compares the envelope of each sentence against all other true envelopes in the corpus rather than against the reconstructed ones.

Because the neural reconstructions preserve the rhythmic properties of the speech envelopes and both axes represent analogous computations of corpus similarity, these two factors combine to produce the strong correlation and gradual change observed in reconstruction accuracies and slopes. If the brain operated as a perfect LTI system yielding nearly flawless envelope reconstructions, the data points would cluster tightly along the diagonal with minimal perpendicular variance. In practice, the scatter plot exhibits residual variance and the estimated slopes are lower than the acoustic similarity between envelopes, which implies that the reconstructions are imperfect. It also shows that despite this imperfection, the acoustic sentence similarity consistently predicted the reconstruction accuracies (slopes) across the three sensor modalities. Rather than treating mismatched sentences as a binary control condition, our results reveal a continuous range of reconstruction accuracies and provide a quantitative measure of how the acoustic structure of a speech corpus influences neural tracking estimates.

#### Temporally Mismatched Envelopes and Reconstructions

To investigate the impact of temporal alignment on reconstruction accuracies between predicted and matched envelopes, we partially mismatched envelope pairs using circular shifts in time to compute a cross-correlation function. These cross-correlation functions were consistent across all three sensor modalities. Notably, the peaks in this neural cross-correlation resembled the peaks computed between all ground truth envelopes using the same circular shift method. Prominent peaks capturing significant reconstruction accuracies and slopes occurred around 1.6 Hz, 1.9 Hz and 4.0 Hz. These frequencies fall into the word and phoneme ranges, broadly aligning with the estimated rates of OLSA words (≈ 2.1 Hz) and syllables (≈ 3.6 Hz), which were also visible in the computed modulation spectrum. This suggests that the neural reconstructions partially captured the linguistic features of the OLSA sentences. Because the quasi-periodic speech envelope reflects linguistic structures at the syllable and word levels (Peelle & Davis, 2012), the periodicities observed in the cross-correlation most likely stem from these linguistic structures. Differences between the neural cross-correlation and the baseline derived from the original envelopes likely reflect imperfect reconstructions (see above) and neural tracking, while the use of circular shifts may have contributed to the slight deviations from the measured word and syllable rates.

#### Null Distributions for Structured Speech

The computation of reconstruction accuracies for mismatched sentences is typically applied to assess the statistical significance of neural tracking by constructing an appropriate null distribution representing chance-level performance. This distribution is usually generated by randomly shuffling the pairings of neural signals and stimulus representations, thereby destroying the true relationship between the brain recordings and the speech signals (Crosse et al., 2021). As a basic control condition (Shuffled Sentences), we randomized trial labels by exchanging sentence envelopes while maintaining their initial temporal alignment. Because OLSA matrix sentences share a rigid syntactic structure, their acoustic envelopes naturally exhibit a mean baseline correlation of ≈ 0.13 (see Figure 6(b)). Consequently, these permuted mismatched envelopes retain a residual correspondence with the brain recordings, yielding to significant reconstruction accuracies. This condition produced an intermediate LMM slope centered among the slopes of individual sentences, which occurs because the envelope reconstructions were compared against a randomized sequence of *all* OLSA envelopes rather than a single specific envelope. The second control condition, which compared the envelope reconstructions to the mean envelope of the OLSA speech material (Mean-Sentence), revealed significant reconstruction accuracies and slopes comparable to those obtained using the true matched envelopes, which quantify neural tracking. This aligns with the high similarity of the mean envelope to all corpus envelopes (≈ 0.32). While it might initially seem surprising that comparing reconstructions to such a generalized template yields such high tracking, this effect is driven by the robust structural similarity shared across matrix sentences. By introducing a temporal shift between reconstructions and envelopes, the cross-correlation revealed dependencies at the word and phoneme levels, indicating that shifts within this range still yielded significant reconstruction accuracies. Applying randomized time shifts entirely breaks this temporal alignment, successfully establishing a strict null distribution centered near zero across all intelligibility levels. This reinforces the importance of this method for generating robust statistical baselines. While using random time shifts is an established and seemingly intuitive procedure, our findings serve as a strong reminder to exercise caution when constructing null distributions for highly structured speech materials.

#### Impact of Initial Sentence Onsets

Due to the sentence onset alignment, the mean envelope (see Figure S2) used for the Mean-Sentence condition revealed a prominent initial peak (≈ 200 ms). To evaluate the degree to which reconstruction accuracies (as measures of neural tracking) were driven by this initial onset response, we recomputed the accuracies and slopes after removing the first 500 ms of each sentence trial. Overall, reconstruction accuracies and fitted slopes decreased for both matched and mismatched sentence envelopes. In particular, reconstruction accuracies declined much more steeply for the mismatched control conditions compared to the matched envelopes. Across all three sensor modalities, LMM slopes decreased to roughly 10 % for the Shuffled Sentences and 50 % for the Mean Sentence control conditions relative to the matched baseline (see Figure S5). This highlights the substantial impact of the aligned initial onset. These findings align with previous research (Chalas et al., 2023; Deoisres et al., 2023), which demonstrated that speech tracking is enhanced around speech onsets. Even without the initial 500 ms, a significant, albeit weaker, correlation remained between the fitted slopes of individual sentences and their acoustic similarity to the OLSA corpus across all three sensor modalities (see Figure S6). However, compared to using the entire sentence, more individual slopes did not differ significantly from zero and a greater number of reconstruction accuracies failed to reach significance relative to their null distributions. Interestingly, the proportion of sentences maintaining significance varied across sensors, decreasing from MEG to EEG and further to ear-EEG. This suggests that whole-head systems, such as MEG and cap-EEG, are more sensitive to the weaker ongoing neural tracking that follows the initial onset response, allowing significant reconstruction accuracies to be derived from the later parts of the sentences.

### Comparing MEG, EEG and ear-EEG

We consistently observed significant reconstruction accuracies and fitted slopes for matched and mismatched sentences across MEG, EEG and ear-EEG, suggesting all three modalities reflect the same underlying neural processes. The strength of these values followed a hierarchy corresponding to spatial coverage (MEG *>* scalp EEG *>* ear-EEG) (Destoky et al., 2019). Although the restricted coverage of ear-EEG reduces its overall sensitivity (Meiser & Bleichner, 2022; Meiser et al., 2020), its physical proximity to the auditory cortex preserves high local sensitivity for speech envelope processing (Brodbeck et al., 2018; Kubanek et al., 2013). Furthermore, previous studies using natural speech demonstrated that ear-EEG performs comparably to cap EEG in objective speech audiometry paradigms (Borges, Alickovic, et al., 2025; Borges, Zaar, et al., 2025a) and auditory attention decoding (Bleichner et al., 2016; Geirnaert et al., 2025; Holtze et al., 2022; Thornton et al., 2024). Quantified numerically using mismatched sentences, the estimated fixed-effect slopes (*β_I_*) (see Figure 5) are highest for MEG (≈ 0.17), slightly lower for EEG (≈ 0.13) and approximately halved for ear-EEG (≈ 0.07), well exceeding the noise floor defined by randomized mismatched speech.

### Effect of Speech-related Features and Speech Material

In this study, we focused exclusively on the speech envelope as a speech-related feature, as it is the most prominent feature in neural tracking research. It would be interesting for future work to investigate how sentence similarity relates to tracking when using more complex stimulus features (such as spectral or cepstral features) or linguistic features like phonetic representations (such as phone probabilities) (Di Liberto et al., 2015). Such features would likely yield different patterns between sentence similarity and tracking, potentially reducing overall dependence due to their higher complexity, which may not be driven by simple mechanisms like onset detection. To overcome the rigid and repetitive structure of matrix test sentences, more ecologically valid natural speech, such as audiobooks, could be utilized. Compared to matrix sentences, natural speech provides greater acoustic variability, enhances neural tracking (Verschueren et al., 2020) and remains highly feasible for objective speech audiometry using EEG or ear-EEG (Borges, Alickovic, et al., 2025; Borges, Zaar, et al., 2025a, 2025b). Observing significant reconstruction accuracies for both matched (heard) and mismatched (unheard) sentences provides further evidence that neural tracking is likely a necessary, but not sufficient, condition for speech perception (Gillis et al., 2022).

## Conclusion

Neural tracking of heard speech yields significant correlations between speech envelopes and neural representations, but comparable correlations can arise for unheard (mismatched) sentences, especially in highly structured speech materials such as the Oldenburg Sentence Test (OLSA). We show that reconstruction accuracies for mismatched envelopes gradually increase with their acoustic similarity to the heard speech material and temporally shifted envelopes reveal peaks reflecting shared phoneme and word rates. Crucially, for mismatched but acoustically similar sentences, these accuracies can reach magnitudes comparable to true envelope tracking. Although correlation magnitudes decreased from MEG to cap-EEG to ear-EEG, the effect remained robust across all three modalities, further supporting the viability of ear-EEG as a portable alternative to whole-head setups.

## Supporting information

Supplemental Material for the article

## Acknowledgments

This work was supported by the Neuroimaging Unit of the Carl von Ossietzky Universität Oldenburg, funded by grants from the German Research Foundation (3T MRI INST 184/152-1 FUGG and MEG INST 184/148-1 FUGG). The authors express their gratitude to all participants for their patience and endurance. We thank Dilek Erdil and Ole Hausendorf for their assistance with data acquisition, Andreas Spiegler for technical support regarding the experimental setup and Daniel Berg for his contributions to the Python implementation of the OLSA framework. We also thank Andreas Spiegler, Stefan Uppenkamp and Thomas Brand for their insightful discussions and feedback throughout the project. During the preparation of this manuscript, the authors used Gemini 3 models (Google) to improve the language and readability of selected sections. Following the use of this AI tool, the authors reviewed and edited the content as necessary and maintain full responsibility for the final version of the manuscript.

## Author contributions

CRediT: **Till Habersetzer:** Conceptualization, Data Curation, Formal Analysis, Investigation, Methodology, Project Administration, Software, Validation, Visualization, Writing – original draft, Writing – review & editing; **Bernd T. Meyer:** Conceptualization, Funding acquisition, Supervision, Visualization, Writing – review & editing; **Andreas Radeloff:** Conceptualization, Funding acquisition, Supervision, Writing – review & editing.

## Data availability statement

The dataset will be made publicly available on OpenNeuro upon publication and will be accompanied by a comprehensive data descriptor article. In the interim, the data supporting the findings of this study are available from the corresponding author upon reasonable request. All analysis scripts are openly accessible via GitHub under the BSD 3 Clause License (https://github.com/tillhabersetzer/neural-tracking-matrix-similarity) and are archived on Zenodo (Habersetzer, 2026).

## Supplemental Material

Supplemental material for this paper is available online.

## References

Akeroyd, M. A., Arlinger, S., Bentler, R. A., Boothroyd, A., Dillier, N., Dreschler, W. A., Gagne, J.-P., Lutman, M., Wouters, J., Wong, L., et al. (2015). International collegium of rehabilitative audiology (icra) recommendations for the construction of multilingual speech tests: Icra working group on multilingual speech tests. International journal of audiology, 54 (sup2), 17–22.

Appelhoff, S., Hurst, A. J., Lawrence, A., Li, A., Mantilla Ramos, Y. J., O’Reilly, C., Xiang, L., Dancker, J., Scheltienne, M., Bialas, O., Alibou, N., Agarwal, A., & Veillette, J. (2025, July). *Pyprep: A python implementation of the preprocessing pipeline (prep) for eeg data.* (Version 0.5.0). Zenodo. 10.5281/zenodo.16039994

Berg, D. (2024, September). *Soundmexpro* (Version 3.1.0.0). Zenodo. 10.5281/zenodo.13847645

Biesmans, W., Das, N., Francart, T., & Bertrand, A. (2016). Auditory-inspired speech envelope extraction methods for improved eeg-based auditory attention detection in a cocktail party scenario. IEEE Trans. Neural Syst. Rehabil. Eng., 25 (5), 402–412.

Bigdely-Shamlo, N., Mullen, T., Kothe, C., Su, K.-M., & Robbins, K. A. (2015). The prep pipeline: Standardized preprocessing for large-scale eeg analysis. Frontiers in neuroinformatics, 9, 16.

Bleichner, M. G., Mirkovic, B., & Debener, S. (2016). Identifying auditory attention with ear-eeg: Ceegrid versus high-density cap-eeg comparison. Journal of neural engineering, 13 (6), 066004.

Borges, H. B., Alickovic, E., Christensen, C. B., Kidmose, P., & Zaar, J. (2025). Age-related differences in eeg-based speech reception threshold estimation using scalp and ear-eeg. Trends in hearing, 29, 23312165251372462.

Borges, H. B., Zaar, J., Alickovic, E., Christensen, C. B., & Kidmose, P. (2025a). The speech reception threshold can be estimated using eeg electrodes in and around the ear. Journal of Neural Engineering, 22 (5), 056008.

Borges, H. B., Zaar, J., Alickovic, E., Christensen, C. B., & Kidmose, P. (2025b). Speech reception threshold estimation via eeg-based continuous speech envelope reconstruction. European Journal of Neuroscience, 61 (6), e70083.

Brand, T., & Wagener, K. (2017). Eigenschaften, leistungen und grenzen von matrixtests. HNO, 65 (3), 182–188.

Brand, T., & Kollmeier, B. (2002). Efficient adaptive procedures for threshold and concurrent slope estimates for psychophysics and speech intelligibility tests. The Journal of the Acoustical Society of America, 111 (6), 2801–2810.

Brodbeck, C., Hong, L. E., & Simon, J. Z. (2018). Rapid transformation from auditory to linguistic representations of continuous speech. Current Biology, 28 (24), 3976–3983.

Brodbeck, C., & Simon, J. Z. (2020). Continuous speech processing. Current Opinion in Physiology, 18, 25–31.

Chalas, N., Daube, C., Kluger, D. S., Abbasi, O., Nitsch, R., & Gross, J. (2023). Speech onsets and sustained speech contribute differentially to delta and theta speech tracking in auditory cortex. Cerebral Cortex, 33 (10), 6273–6281.

Cooper, J. K., Vanthornhout, J., van Wieringen, A., & Francart, T. (2025). Objectively measuring audiovisual effects in noise using virtual human speakers. Trends in Hearing, 29, 23312165251333528.

Crosse, M. J., Di Liberto, G. M., Bednar, A., & Lalor, E. C. (2016). The multivariate temporal response function (mtrf) toolbox: A matlab toolbox for relating neural signals to continuous stimuli. Frontiers in human neuroscience, 10, 604.

Crosse, M. J., Zuk, N. J., Di Liberto, G. M., Nidiffer, A. R., Molholm, S., & Lalor, E. C. (2021). Linear modeling of neurophysiological responses to speech and other continuous stimuli: Methodological considerations for applied research. Frontiers in neuroscience, 15, 705621.

Deoisres, S., Lu, Y., Vanheusden, F. J., Bell, S. L., & Simpson, D. M. (2023). Continuous speech with pauses inserted between words increases cortical tracking of speech envelope. PloS One, 18 (7), e0289288.

Destoky, F., Philippe, M., Bertels, J., Verhasselt, M., Coquelet, N., Vander Ghinst, M., Wens, V., De Tiège, X., & Bourguignon, M. (2019). Comparing the potential of meg and eeg to uncover brain tracking of speech temporal envelope. Neuroimage, 184, 201–213.

Di Liberto, G. M., O’Sullivan, J. A., & Lalor, E. C. (2015). Low-frequency cortical entrainment to speech reflects phoneme-level processing. Current Biology, 25 (19), 2457–2465.

Ding, N., & Simon, J. Z. (2013). Adaptive temporal encoding leads to a background-insensitive cortical representation of speech. Journal of Neuroscience, 33 (13), 5728–5735.

Elliott, T. M., & Theunissen, F. E. (2009). The modulation transfer function for speech intelligibility. PLoS computational biology, 5 (3), e1000302.

Etard, O., & Reichenbach, T. (2019). Neural speech tracking in the theta and in the delta frequency band differentially encode clarity and comprehension of speech in noise. Journal of Neuroscience, 39 (29), 5750–5759.

Geirnaert, S., Kappel, S. L., & Kidmose, P. (2025). A direct comparison of simultaneously recorded scalp, around-ear and in-ear eeg for neural selective auditory attention decoding to speech. Scientific Reports, 15 (1), 41655.

Gillis, M., Van Canneyt, J., Francart, T., & Vanthornhout, J. (2022). Neural tracking as a diagnostic tool to assess the auditory pathway. Hearing Research, 426, 108607.

Gramfort, A., Luessi, M., Larson, E., Engemann, D. A., Strohmeier, D., Brodbeck, C., Goj, R., Jas, M., Brooks, T., Parkkonen, L., et al. (2013). Meg and eeg data analysis with mne-python. Frontiers in Neuroinformatics, 7, 267.

Habersetzer, T. (2026, September). *Code for: Analysis of the influence of gradual changes in matrix sentence similarity on neural envelope tracking* (Version v1.0.0). Zenodo. 10.5281/zenodo.22692704

Habersetzer, T., Radeloff, A., & Meyer, B. T. (2026). Comparing neurophysiological correlates of speech-in-noise perception across meg, eeg and ear-eeg. bioRxiv. 10.64898/2026.07.31.741889

Habersetzer, T., Steuer, S., Radeloff, A., & Meyer, B. T. (2025). Meg-scans-a comprehensive magnetoencephalography speech dataset with stories, chirps and noisy sentences. Scientific Data.

Hagerman, B. (1982). Sentences for testing speech intelligibility in noise. Scandinavian audiology, 11 (2), 79–87.

Holtze, B., Rosenkranz, M., Jaeger, M., Debener, S., & Mirkovic, B. (2022). Ear-eeg measures of auditory attention to continuous speech. Frontiers in Neuroscience, 16, 869426.

Humes, L. E. (2019). The world health organization’s hearing-impairment grading system: An evaluation for unaided communication in age-related hearing loss. International journal of audiology, 58 (1), 12–20.

Karunathilake, I. D., Dunlap, J. L., Perera, J., Presacco, A., Decruy, L., Anderson, S., Kuchinsky, S. E., & Simon, J. Z. (2023). Effects of aging on cortical representations of continuous speech. Journal of neurophysiology, 129 (6), 1359–1377.

Kollmeier, B., Warzybok, A., Hochmuth, S., Zokoll, M. A., Uslar, V., Brand, T., & Wagener, K. C. (2015). The multilingual matrix test: Principles, applications, and comparison across languages: A review. International journal of audiology, 54 (sup2), 3–16.

Kubanek, J., Brunner, P., Gunduz, A., Poeppel, D., & Schalk, G. (2013). The tracking of speech envelope in the human cortex. PloS one, 8 (1), e53398.

Lesenfants, D., Vanthornhout, J., Verschueren, E., Decruy, L., & Francart, T. (2019). Predicting individual speech intelligibility from the cortical tracking of acoustic-and phonetic-level speech representations. Hearing research, 380, 1–9.

Meiser, A., & Bleichner, M. G. (2022). Ear-eeg compares well to cap-eeg in recording auditory erps: A quantification of signal loss. Journal of Neural Engineering, 19 (2), 026042.

Meiser, A., Tadel, F., Debener, S., & Bleichner, M. G. (2020). The sensitivity of ear-eeg: Evaluating the source-sensor relationship using forward modeling. Brain topography, 33 (6), 665–676.

Muncke, J., Kuruvila, I., & Hoppe, U. (2022). Prediction of speech intelligibility by means of eeg responses to sentences in noise. Frontiers in Neuroscience, 16, 876421.

Nuesse, T., Wiercinski, B., Brand, T., & Holube, I. (2019). Measuring speech recognition with a matrix test using synthetic speech. Trends in Hearing, 23, 2331216519862982.

Obleser, J., & Kayser, C. (2019). Neural entrainment and attentional selection in the listening brain. Trends in cognitive sciences, 23 (11), 913–926.

Oostenveld, R., Fries, P., Maris, E., & Schoffelen, J.-M. (2011). Fieldtrip: Open source software for advanced analysis of meg, eeg, and invasive electrophysiological data. Computational intelligence and neuroscience, 2011 (1), 156869.

Peelle, J. E., & Davis, M. H. (2012). Neural oscillations carry speech rhythm through to comprehension. Frontiers in psychology, 3, 320.

Peirce, J., Gray, J. R., Simpson, S., MacAskill, M., Höchenberger, R., Sogo, H., Kastman, E., & Lindeløv, J. K. (2019). Psychopy2: Experiments in behavior made easy. Behavior research methods, 51 (1), 195–203.

Ratelle, D., & Tremblay, P. (2025). Neural tracking of continuous speech in adverse acoustic conditions among healthy adults with normal hearing and hearing loss: A systematic review. Hearing Research, 109367.

Shannon, R. V., Zeng, F.-G., Kamath, V., Wygonski, J., & Ekelid, M. (1995). Speech recognition with primarily temporal cues. Science, 270 (5234), 303–304.

Taulu, S., & Kajola, M. (2005). Presentation of electromagnetic multichannel data: The signal space separation method. Journal of Applied Physics, 97 (12).

Taulu, S., & Simola, J. (2006). Spatiotemporal signal space separation method for rejecting nearby interference in meg measurements. Physics in Medicine & Biology, 51 (7), 1759.

Thornton, M., Mandic, D., & Reichenbach, T. (2024). Comparison of linear and nonlinear methods for decoding selective attention to speech from ear-eeg recordings. arXiv preprint arXiv:2401.05187.

Vanthornhout, J., Decruy, L., Wouters, J., Simon, J. Z., & Francart, T. (2018). Speech intelligibility predicted from neural entrainment of the speech envelope. Journal of the Association for Research in Otolaryngology, 19 (2), 181–191.

Verschueren, E., Vanthornhout, J., & Francart, T. (2020). The effect of stimulus choice on an eeg-based objective measure of speech intelligibility. Ear and hearing, 41 (6), 1586–1597.

Wagener, K., Brand, T., & Kollmeier, B. (1999a). Entwicklung und Evaluation eines Satztests für die deutsche Sprache Teil II: Optimierung des Oldenburger Satztests (Development and evaluation of a German speech intelligibility test. Part II: Optimization of the Oldenburg sentence test). Zeitschrift für Audiologie, 38, 44–56.

Wagener, K., Brand, T., & Kollmeier, B. (1999b). Entwicklung und Evaluation eines Satztests für die deutsche Sprache Teil III: Evaluation des Oldenburger Satztests (Development and evaluation of a German speech intelligibility test. Part III: Evaluation of the Oldenburg sentence test). Zeitschrift für Audiologie, 38, 86–95.

Wagener, K., Kühnel, V., & Kollmeier, B. (1999). Entwicklung und Evaluation eines Satztests für die deutsche Sprache I: Design des Oldenburger Satztests (Development and evaluation of a German speech intelligibility test. Part I: Design of the Oldenburg sentence test). Zeitschrift für Audiologie, 38, 4–15.

Widmann, A., Schröger, E., & Maess, B. (2015). Digital filter design for electrophysiological data–a practical approach. Journal of neuroscience methods, 250, 34–46.

