## Supplemental Material for the article for "Analysis of the influence of gradual changes in matrix sentence similarity on neural envelope tracking"

#### Contents

**Overview** **2**

**Layout EEG-Cap** **2**

**Subject-Specific Exclusions and Protocol Deviations** **2**

**Technical Details Concerning FDR Correction** **3**

**Mean-Sentence Envelope** **4**

**Analysis Examples for Reconstruction Accuracies with Sentence Similarity and Time-Shift** **4**

**Robustness Check: Onset Response Removed** **6**

Reconstruction Accuracies Across Control Conditions . . . . . . . . . . 6

Reconstruction Accuracies and Sentence Similarity . . . . . . . . . . 6

### Overview

This supplemental document provides details on the layout of the costum EEG-cap, subject-specific exclusions and protocol deviations, FDR computations and a depiction of the mean envelope used for the Mean-Sentence control condition. Additionally, it contains examples of computed reconstruction accuracies and fitted slopes for selected OLSA sentences and shift latencies between reconstructions and matched sentence envelopes. Finally, it presents the reconstruction accuracies and neural tracking measures obtained after removing the first 500 ms of the reconstructions to exclude the initial onset response for each sentence.

### Layout EEG-Cap

Figure S1(a) illustrates the layout of the custom 76-channel EEG-cap used during the recordings (BC-MEG-76-X2; Easycap, Wörthsee, Germany). The layout and channel assignments correspond to the standard Elekta Neuromag 64-channel EEG-cap, with the addition of 12 around-ear electrodes. The ear-EEG subset defined for this analysis consists of these 12 around-ear electrodes along with standard cap channels T7, TP7, T8 and TP8, totaling 16 ear-EEG channels (8 channels per ear). The right-hemisphere portion of this subset is highlighted by a shaded gray area in Figure S1(b).

**(a)** **(b)**

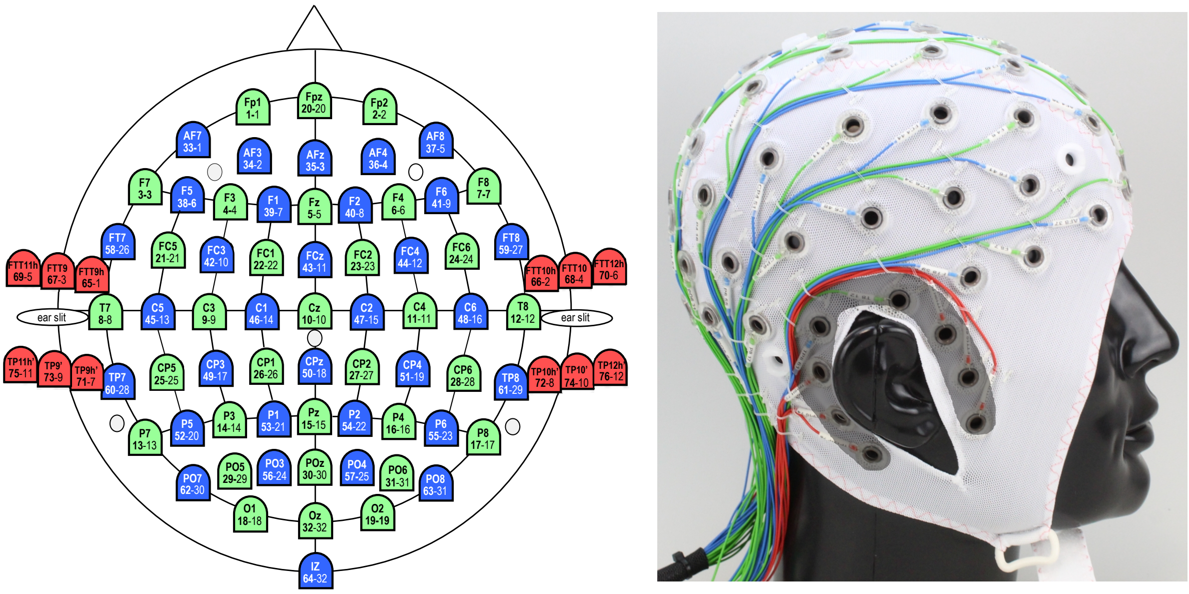


Figure S1: Custom 76-channel EEG-cap layout and ear-EEG subset. **(a)** Layout of the custom EEG-cap (BC-MEG-76-X2) used during the recordings. The standard 64-channel layout comprises the green and blue electrodes, while the 12 additional around-ear electrodes are shown in red. **(b)** Photograph of the cap worn on a dummy head, showing the right side. The eight electrodes that make up the right-hemisphere ear-EEG subset are highlighted by a shaded gray area. Both images were modified and reprinted with permission of Easycap GmbH.

### Subject-Specific Exclusions and Protocol Deviations

The finalized experimental design, comprising three sessions, continuous Head Position Tracking (cHPI) and individualized Speech Reception Threshold (SRT) values, was fully implemented starting with subject sub-02 (see Table S1). Data from subjects sub-00 (pilot) and sub-01 were included in the analysis despite the following protocol deviations. For subject sub-00, a preliminary version of the paradigm utilized a fixed SRT (−9.6 dB, slope: 0.13) rather than individualized values and cHPI was inactive. Furthermore, this pilot session included only five signal-to-noise-ratios (SNR) levels: the −40 dB condition was omitted and a clean speech condition (no noise) was used instead of the 0 dB SNR condition. Subject sub-01 followed the finalized paradigm regarding individualized SRTs and active cHPI. However, the lowest SNR was set to −20 dB instead of the −40 dB level used for all subsequent participants. Both subjects, sub-00 and sub-01, followed a two-session design, while the finalized protocol comprised three sessions. Consequently, for these initial two subjects, screening was integrated into the first MEG/EEG recording session (precluding separate behavioral OLSA screening data), thereby limiting the training phase to one OLSA list instead of three.

Table S1: Protocol deviations for early subjects compared to the final experimental design.

| **Feature** | **sub-00 (Pilot)** | **sub-01** | **sub-02 onwards** |
|---|---|---|---|
| SRT | Fixed (−9.6 dB) | Individualized | Individualized |
| Highest SNR Cond. | Clean Speech | 0 dB | 0 dB |
| Lowest SNR Cond. | N/A (5 conditions) | −20 dB | −40 dB |
| Total Sessions | 2 | 2 | 3 |
| Training Lists | 1 | 1 | 3 |
| cHPI | No | Yes | Yes |
| Screening | Mixed in Session 1 | Mixed in Session 1 | Separate Session |

All subjects were included in the final analysis. Subjects sub-01 and sub-19 were retained despite slightly elevated hearing thresholds in one ear, as their inclusion did not alter the overall results. For consistency across the dataset, the clean speech condition for sub-00 was remapped to 0 dB. Furthermore, the −40 dB (−20 dB for sub-01) and 0 dB conditions were remapped to 0 % and 100 % intelligibility for all subjects, respectively. This mapping reflects behavioral performance that was consistently at floor (≤1 %) or ceiling (≥98 %) at these levels.

### Technical Details Concerning FDR Correction

To control the False Discovery Rate (FDR), the Benjamini-Hochberg procedure was applied throughout the analysis. Table S2 summarizes which statistical tests had their p-values corrected, the number of simultaneous tests ($n_{tests}$), and the underlying combinatorial calculations for each figure.

Table S2: Summary of FDR corrections across figures.

| **Figure** | **Statistical Test** | $n_{tests}$ | **Derivation / Combinatorics** |
|---|---|---|---|
| Figure 3 | Pearson correlation ($\rho$) | 3 | 3 sensors |
| | Fixed-effect slope ($\beta_I$) | 3 | 3 sensors |
| Figure 5 | Pearson correlation ($\rho$) | 3 | 3 sensors |
| | Null distribution comparison | 1800 | 3 sensors × 6 SNRs × 100 sentences |
| | Fixed-effect slope ($\beta_I$) | 300 | 3 sensors × 100 sentences |
| Figure 6 | Null distribution comparison | 2322 | 3 sensors × 6 SNRs × 129 shifts |
| | Fixed-effect slope ($\beta_I$) | 387 | 3 sensors × 129 shifts |

### Mean-Sentence Envelope

The Mean-Sentence envelope, calculated as the average of all 100 OLSA sentence envelopes, is depicted in Figure S2.

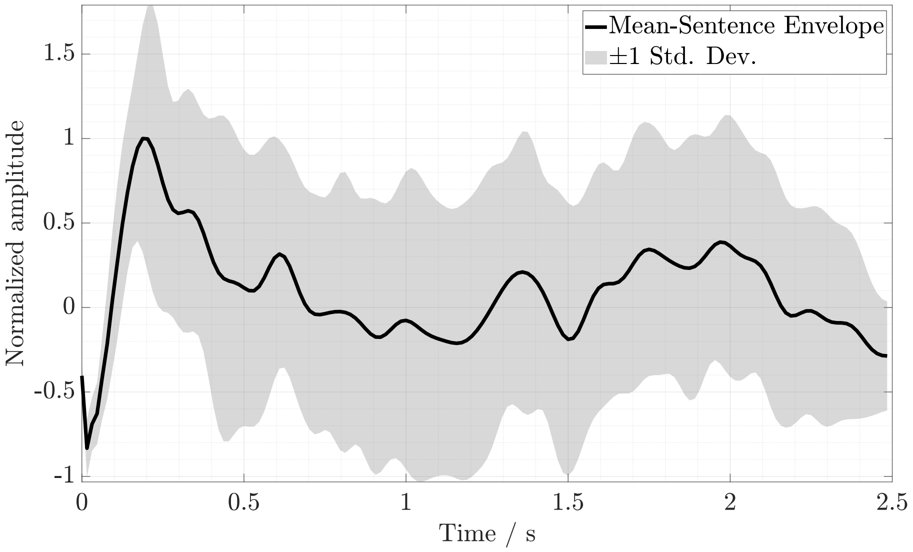


Figure S2: The Mean-Sentence envelope (black line) shown alongside $\pm 1$ standard deviation of the individual averaged OLSA envelopes. For visualization purposes, the signals have been normalized using peak amplitude scaling so that the mean envelope lies within a range of $[-1, 1]$.

### Analysis Examples for Reconstruction Accuracies with Sentence Similarity and Time-Shift

During the primary analysis, reconstruction accuracies and fitted slopes were computed for various OLSA sentences and circular time-shifts. Figure S3 illustrates the reconstruction accuracies and fitted linear mixed-effects models (LMMs) for three out of the 100 OLSA sentences. For each sentence, its corresponding envelopes were concatenated and compared with the decoder's reconstruction. This approach yielded a distinct fitted model for each sentence, revealing that the resulting slope correlates with the acoustic similarity between the individual sentence envelope and the broader OLSA speech material.

In a subsequent analysis, Figure S4 presents the reconstruction accuracies and fitted linear mixed-effects models (LMMs) for three selected time-shifts between the reconstructed and matched sentence envelopes. As demonstrated, applying different circular time-shifts systematically led to variations in reconstruction accuracies and their corresponding fitted slopes.

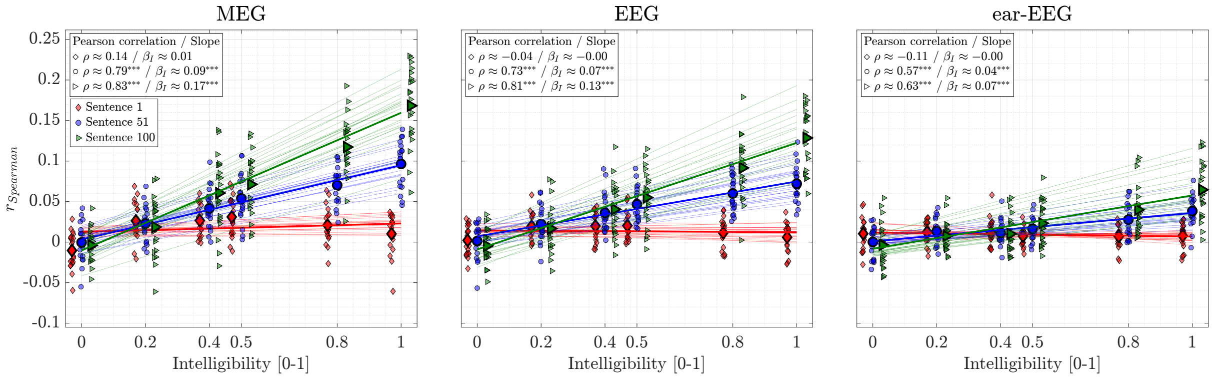


Figure S3: Reconstruction accuracy ($r_{Spearman}$) as a function of speech intelligibility for MEG, EEG and ear-EEG. Individual data points are shown alongside grand average means (large symbols) for three selected OLSA sentences (sentences 1, 51 and 100). These sentences are numbered in ascending order based on their mean acoustic similarity to the OLSA speech material, meaning sentence 100 exhibits the highest similarity. To improve visibility, x-axis positions included a horizontal offset between conditions and a slight jitter for individual data points. Thick lines represent LMM fixed-effects, while thin lines represent individual fits. Inset text displays Pearson correlation coefficients ($\rho$) between intelligibility and $r_{Spearman}$, alongside fixed-effect slope estimates ($\beta_I$). Intelligibility is scaled as a proportion (0–1), yielding a dimensionless slope. All $p$-values were adjusted using FDR. Asterisks indicate significance: *** $p < 0.001$, ** $p < 0.01$, * $p < 0.05$.

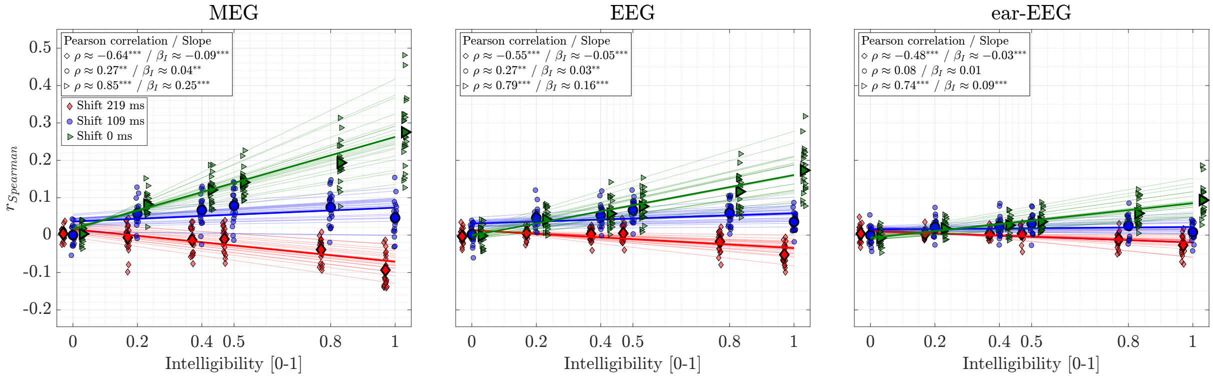


Figure S4: Reconstruction accuracy ($r_{Spearman}$) as a function of speech intelligibility for MEG, EEG and ear-EEG. Individual data points are shown alongside grand average means (large symbols) for three selected time-shifts between the reconstructed and matched sentence envelopes (0 ms, 109 ms and 219 ms). To improve visibility, x-axis positions included a horizontal offset between conditions and a slight jitter for individual data points. Thick lines represent LMM fixed-effects, while thin lines represent individual fits. Inset text displays Pearson correlation coefficients ($\rho$) between intelligibility and $r_{Spearman}$, alongside fixed-effect slope estimates ($\beta_I$). Intelligibility is scaled as a proportion (0–1), yielding a dimensionless slope. All $p$-values were adjusted using FDR. Asterisks indicate significance: *** $p < 0.001$, ** $p < 0.01$, * $p < 0.05$.

### Robustness Check: Onset Response Removed

In the following results, the first 500 ms of the neurophysiological responses (MEG, EEG and ear-EEG) were removed for each OLSA sentence to eliminate the initial stimulus-onset response. Consequently, only the remaining segments of the sentence envelopes (> 500 ms) were used to compute both, the reconstruction accuracies and the acoustic similarities between envelope pairs.

#### Reconstruction Accuracies Across Control Conditions

Figure S5 displays reconstruction accuracies and fitted LMMs across the four control conditions. Compared to the analysis using the complete response that included the onset response (see Figure 3 in publication), the reconstruction accuracies ($r_{Spearman}$), the fitted slopes ($\beta_I$) and the Pearson correlation coefficients ($\rho$) are lower in all three sensor modalities. Nonetheless, the Shuffled Sentences and Mean-Sentence conditions still exhibit an increase in reconstruction accuracy with speech intelligibility. In contrast to Figure 3, the Mean-Sentence condition shows reduced reconstruction accuracies and a reduced slope compared to the Matched Sentences condition.

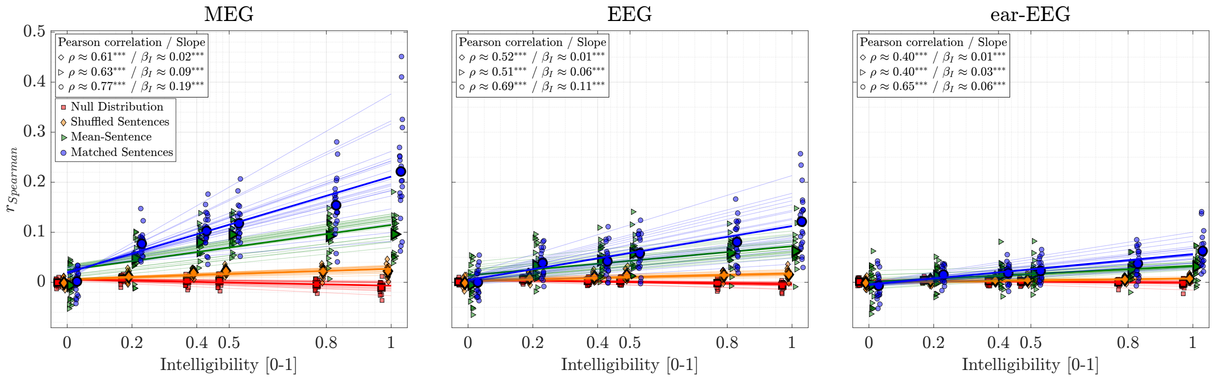


Figure S5: Reconstruction accuracy ($r_{Spearman}$) as a function of speech intelligibility for MEG, EEG and ear-EEG (onset response removed). Individual data points are shown alongside grand average means (large symbols) for four conditions: Matched Sentences (blue circles), Mean-Sentence (green triangles), Shuffled Sentences (orange diamonds) and Null Distribution (red squares). To improve visibility, x-axis positions included a horizontal offset between conditions and a slight jitter for individual data points. Thick lines represent LMM fixed-effects, while thin lines represent individual fits. Inset text displays Pearson correlation coefficients ($\rho$) between intelligibility and $r_{Spearman}$, alongside fixed-effect slope estimates ($\beta_I$). Intelligibility is scaled as a proportion (0–1), yielding a dimensionless slope. All $p$-values were adjusted using FDR. Asterisks indicate significance: *** $p < 0.001$, ** $p < 0.01$, * $p < 0.05$.

#### Reconstruction Accuracies and Sentence Similarity

The fixed-effect slopes ($\beta_I$), computed for concatenated identical OLSA sentence envelopes with the onset removed, are plotted against their corresponding mean acoustic similarity scores ($\rho_{mean}$) in Figure S6. All three sensor modalities exhibit a significant Pearson correlation between the fitted slopes and acoustic similarity, even when the onset response is excluded. Compared to Figure 5 in the publication, the fixed-effect slopes, acoustic similarities between envelopes and overall Pearson correlations are reduced, highlighting the substantial impact of the onset response. Nevertheless, the main effects remain robust. However, these reduced values result in a greater number of fitted slopes that do not significantly differ from zero, as well as more sentences failing to show a significant difference in reconstruction accuracy from the null distribution.

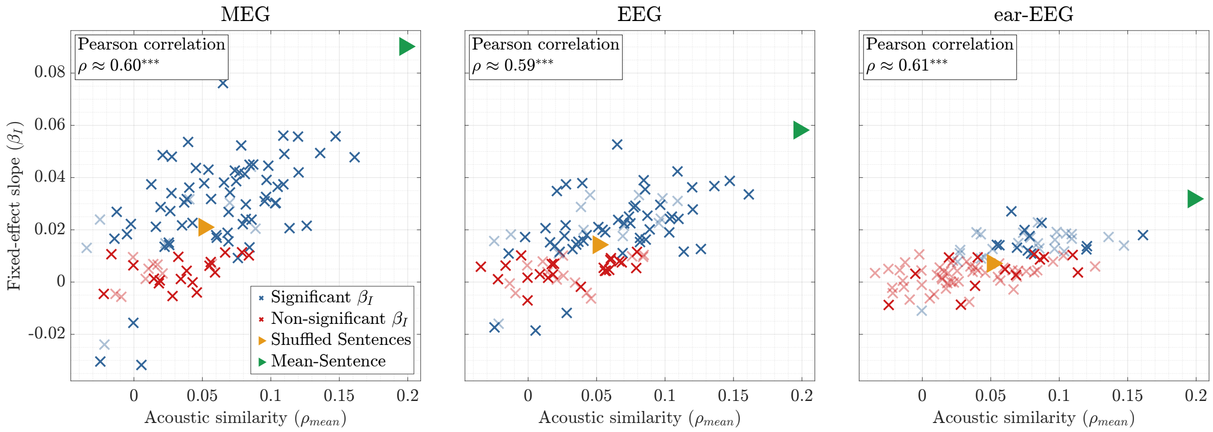


Figure S6: Relationship between sentence-specific mean acoustic similarity and fitted slope (onset response removed). Fixed-effect slopes ($\beta_I$) are plotted against mean acoustic envelope similarity ($\rho_{mean}$) for individual OLSA sentences (represented by crosses). Blue crosses indicate sentences with a significant fixed-effect slope (different from zero), while red crosses indicate non-significant slopes. Sentences whose reconstruction accuracies did not show a significant difference from their null distributions are marked with faded crosses. Pearson correlation coefficients ($\rho$) between $\beta_I$ and $\rho_{mean}$ are provided for MEG, EEG and ear-EEG. Correlations were calculated across all sentences, excluding the Mean-Sentence and Shuffled Sentences conditions. For reference, Mean-Sentence (green triangle) and Shuffled Sentences (orange triangle) conditions are indicated. Asterisks denote significant Pearson correlations (*** $p < 0.001$, ** $p < 0.01$, * $p < 0.05$). All tests were FDR corrected.
